# Quick and robust method for the generation of human iPSC-derived choroid plexus organoids

**DOI:** 10.1101/2025.08.20.671243

**Authors:** Rodi Kado Abdalkader, Takuya Fujita

## Abstract

The choroid plexus (ChP) is a key brain structure responsible for cerebrospinal fluid (CSF) production and forms a selective barrier that regulates brain homeostasis and immune surveillance. In vitro models of ChP are essential for studying CSF dynamics, viral entry, neuroinflammation, and CNS drug transport, yet current organoid protocols remain complex, slow, and difficult to reproduce.

Here, we report a quick and robust method for the generation of human iPSC-derived ChP organoids that is xeno-free/ serum-free, scalable, and reproducible. Early GSK3β inhibition and transient WNT modulation guide organoids toward cystic ChP-enriched structures, confirmed by ventricle-like morphology and expression of canonical markers (TTR, ZO-1).

This minimal workflow enables rapid production of functional ChP-like organoids suitable for CSF physiology studies, barrier modeling, and translational neuroscience, providing an accessible platform for both basic and applied research.

## 1. Introduction

The choroid plexus (ChP) plays an essential role in brain development and homeostasis by producing cerebrospinal fluid (CSF) and forming the blood–CSF barrier[1,2]. It has emerged as a critical gateway for neuroinflammation, viral infections (e.g., Zika, HIV and SARS-CoV-2)[3–5], and drug transport[5–7], making it an attractive target for translational research.

Despite its importance, in vitro modelling of the ChP remains challenging. Current protocols for ChP organoids [8,9]require long culture times, complex stepwise signalling, or serum supplementation, which limits scalability, reproducibility, and adoption.

Here, we present a quick and robust method for generating human induced pluripotent stem cell (iPSC)-derived ChP organoids that is minimal, xeno-free/ serum-free, and reproducible. Our workflow rapidly produces cystic organoids enriched for ChP epithelium, expressing canonical markers and forming ventricle-like structures, thus providing a practical platform for CSF physiology, neuroinflammation modelling, and CNS drug testing.

## 2. Materials and methods

### 2.1. Human iPSC culture

Human iPSC line 201B7-Ff (Riken Cell Bank, Japan)[10] was cultured under protocols approved by the Ritsumeikan University Ethics Committee (No. 2021-004-3). Cells were maintained on iMatrix-511–coated plates in mTeSR™ Plus medium (STEMCELL Technologies), with 10 µM Y-27632 during initial seeding (1×10^4^ cells/well in 6-well plates). Medium was refreshed every 2 days. For passaging, cells were dissociated with 0.5**×** TrypLE Select (Thermo Fisher Scientific), centrifuged at 160 ×g for 5 min, and replated in mTeSR™ Plus with Y-27632.

### 2.2. Organoid generation

For organoid induction, single-cell suspensions were seeded into low-adhesion 96-well plates (1×10^5^ cells/well) in mTeSR™ Plus with Y-27632 to form spheroids within 24 h. Spheroids were exposed to the GSK3β inhibitor CHIR99021 from days 0–3, either alone or with the TGFβ receptor inhibitor SB505124 in E6 medium. From days 3–6, WNT signalling was transiently inhibited using IWP2 (Supplementary Table 1). Selected spheroids were embedded in Matrigel (44:1 v/v) (Thermo Fisher Scientific) during this period to promote structural support. Retinoic acid was added from days 6–12 to enhance dorsal forebrain and ChP specification. Medium changes were performed every 2–3 days.

### 2.3. Histology and immunofluorescence

Organoids were fixed in 4% PFA, paraffin-embedded, and sectioned (5 µm). H&E staining followed standard protocols. For immunofluorescence, sections underwent Tris-EDTA (pH 9.0) antigen retrieval, 0.5% Triton X-100 permeabilization, and blocking in 5% BSA/0.1% Tween-20. Samples were incubated overnight at 4°C with primary antibodies against TTR, ZO-1, PAX6, Tuj-1, and GFAP (Supplementary Table 2), followed by Alexa Fluor 488 or 555 secondary antibodies (1:500; Thermo Fisher Scientific) and DAPI nuclear counterstain. Imaging was performed with a Keyence fluorescence microscope.

### 2.4. Statistical analysis and data visualization

All experiments were performed with ≥3 biological replicates. Data were analyzed using GraphPad Prism 8 with Dunnett’s, paired t-test, or Tukey’s tests as appropriate. Graphs were generated using GraphPad Prism 8.

## 3. Results

We developed a xeno-free/ serum-free rapid method for generating human iPSC-derived ChP organoids by modulating early GSK3β and WNT signaling. As illustrated in the experimental timeline (Figure 1a), four culture conditions (ORG_1–ORG_4) were tested, with combinations of CHIR99021, SB505124, transient IWP2 treatment, and Matrigel embedding. By day 6, spheroids from all conditions formed uniformly, though ORG_3 and ORG_4 began to show signs of surface irregularities indicative of cyst initiation (Figure 1b). By day 20, ORG_1 and ORG_2 organoids retained a compact neuroepithelial morphology, while ORG_3 and ORG_4 developed large cystic structures (Figure 2a). Quantification confirmed a significant increase in organoid area in the cystic groups (ORG_3 and ORG_4) compared to compact groups (Figure 2b). Furthermore, cyst formation was enhanced under Matrigel embedding (ORG_4), showing distinct lumens (Figure 2c). Histological (H&E) and immunofluorescence analyses on day 30 revealed clear regional identity differences between groups. ORG_1 and ORG_2 organoids exhibited dorsal forebrain-like neuroepithelial layering, with widespread expression of PAX6 and Tuj1, markers of early cortical development (Figure 3a). In contrast, ORG_4 organoids displayed large, ventricle-like cavities lined with a monolayer of epithelial protrusions, a morphology resembling choroid plexus villi, as revealed by H&E staining. These structures were further supported by immunofluorescence showing expression of canonical ChP markers TTR and ZO-1, along with scattered GFAP^+^ astrocytes (Figure 3b). Quantitative analysis revealed that Matrigel embedding significantly increased the proportion of Tuj1^+^ cells and reduced PAX6^+^ populations in ORG_1 (Figure 3c), supporting enhanced neuronal maturation. In ORG_4, approximately 20% of the epithelial cells were TTR-positive, while a distinct proportion remained Tuj1^+^, indicating a mixed population of ChP and neuronal lineage cells (Figure 3d). Together, these findings confirm that early GSK3β/WNT modulation combined with Matrigel embedding reliably generates cystic, ChP-enriched organoids within a rapid and minimal culture framework.

**Figure 1.**
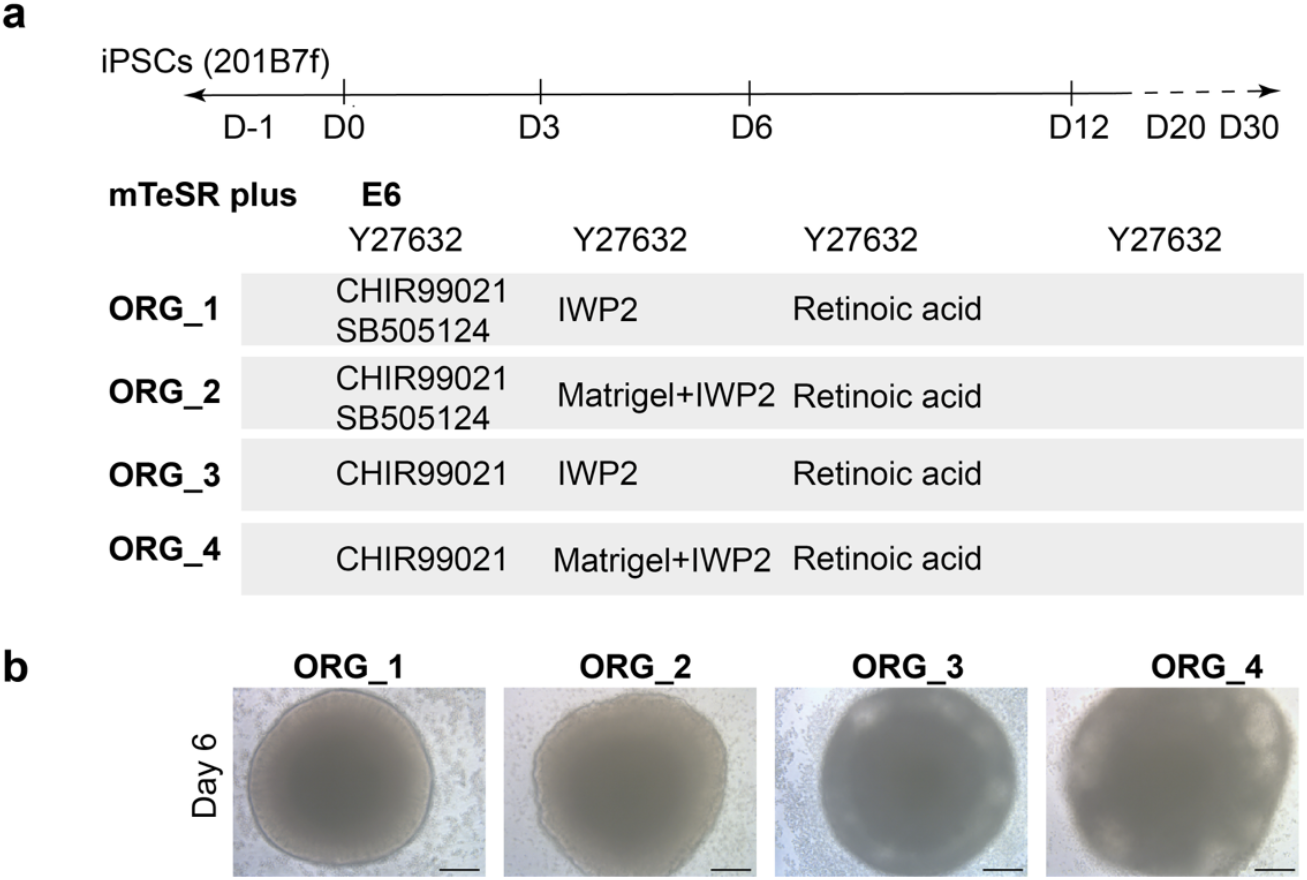
Experimental timeline and morphology of organoids under different early signaling conditions. **(a)** Schematic timeline of organoid induction from human iPSCs (201B7f), showing key medium changes and signaling modulation steps from day −1 to day 30. Four experimental groups (ORG_1 to ORG_4) were exposed to CHIR99021 with or without SB505124 from days 0 to 3, followed by WNT inhibition using IWP2, with or without Matrigel embedding between days 3 and 6. Retinoic acid was added from day 6 to day 12 to promote neural specification. **(b)** Representative brightfield images of organoids at day 6 from each group (ORG_1 to ORG_4), showing differences in morphology and size during early stages of development. Scale bars: 200 µm.

**Figure 2.**
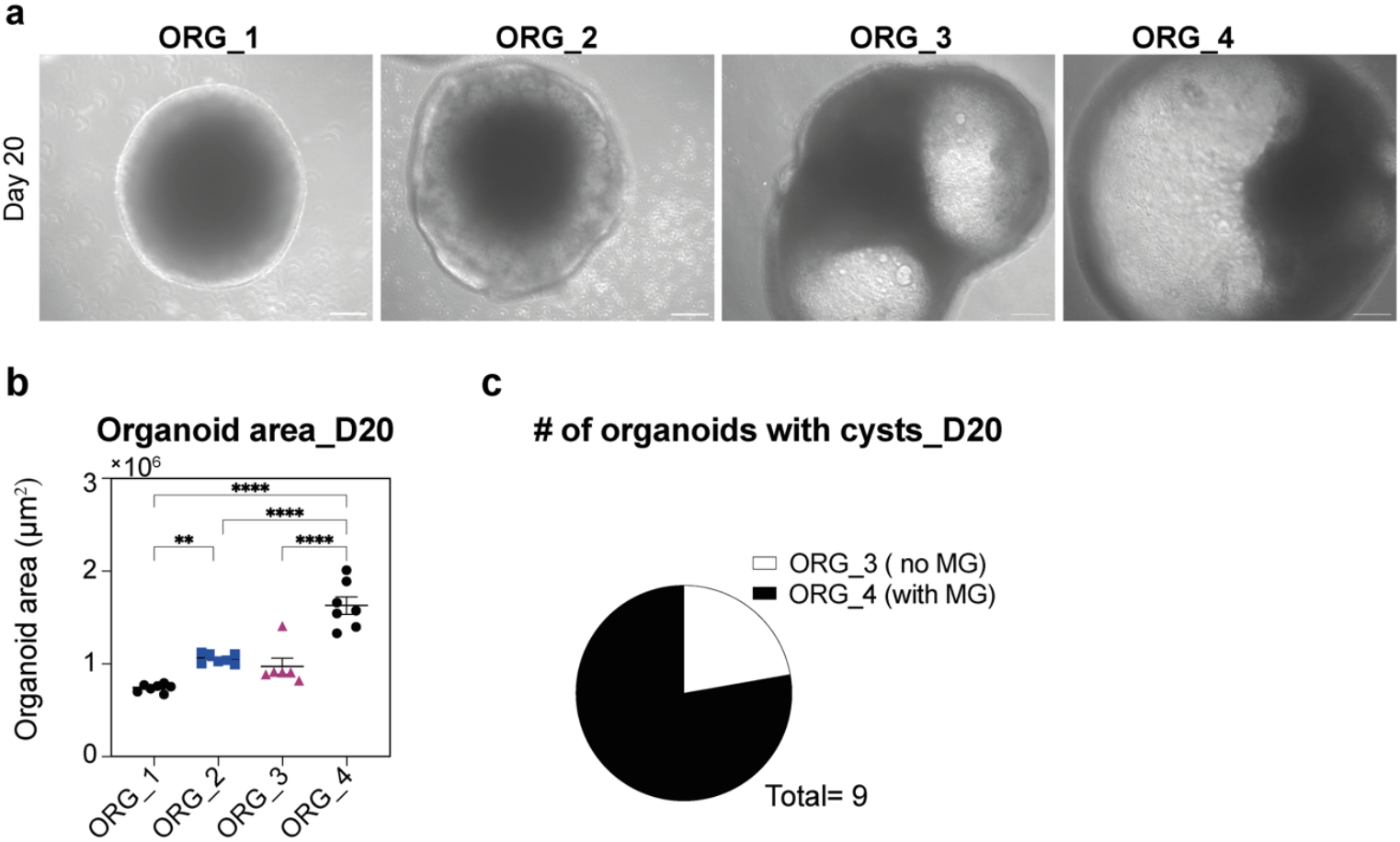
Early GSK3β and WNT modulation regulates organoid morphology and cyst formation. **(a)** Brightfield images of organoids at day 20 from each treatment group. Organoids exposed to CHIR99021 and SB505124 (ORG_1, ORG_2) maintained a compact morphology, while those treated with CHIR99021 alone (ORG_3, ORG_4) exhibited prominent cyst formation, particularly under Matrigel embedding conditions (ORG_4). Scale bars: 200 µm. **(b)** Quantification of organoid area at day 20. Cystic organoids (ORG_3, ORG_4) showed significantly larger areas compared to compact organoids (ORG_1, ORG_2). Data represent mean ± SEM. Statistical significance determined by one-way ANOVA with Tukey’s post hoc test; ^**^p < 0.01, ^**^p < 0.0001. **(c)** Pie chart showing the number of organoids with visible cysts in ORG_3 (no Matrigel) and ORG_4 (with Matrigel).

**Figure 3.**
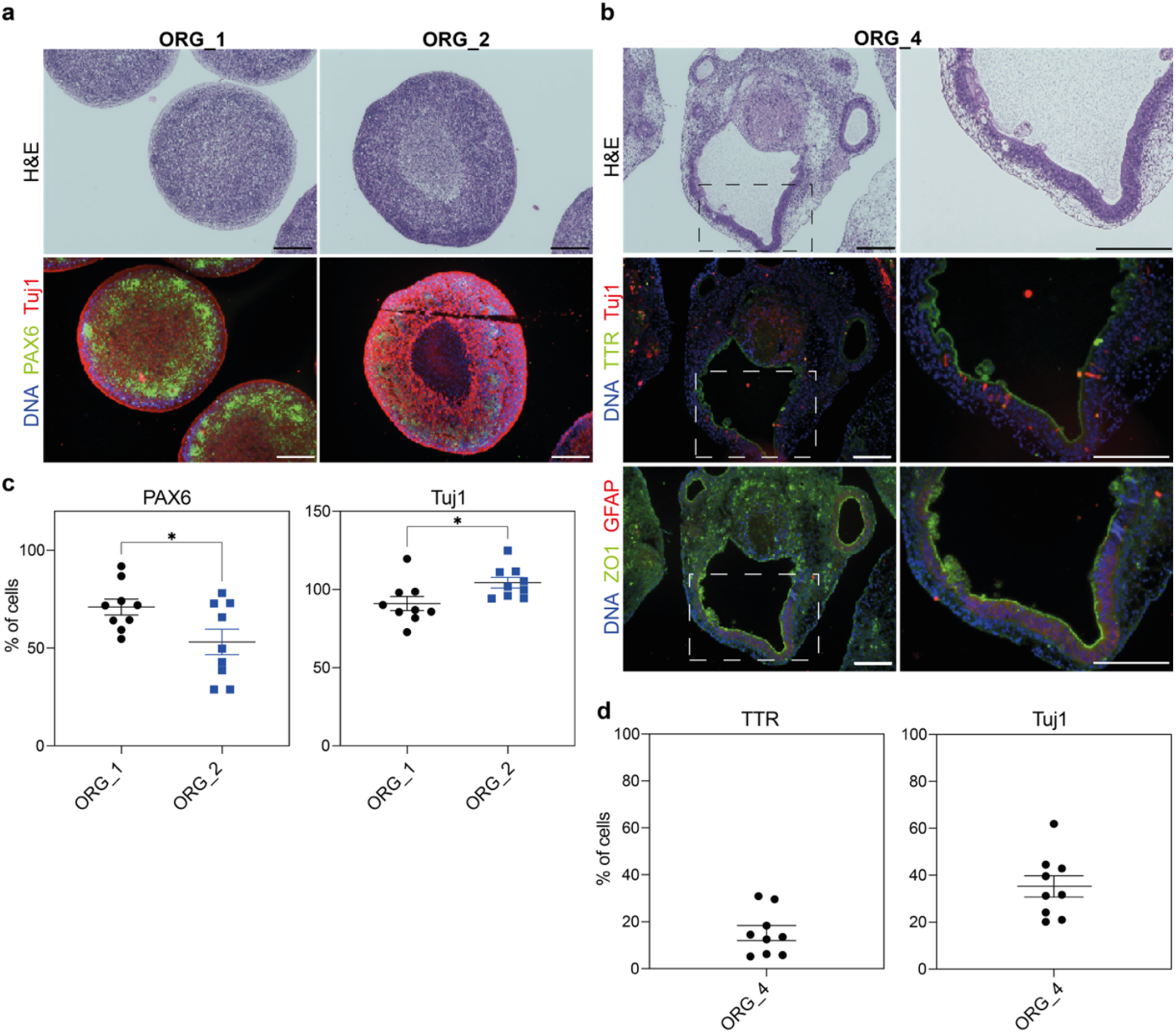
Histological and molecular characterization confirms dorsal forebrain and choroid plexus identities. **(a)** Representative H&E and immunofluorescence images of ORG_1 and ORG_2 organoids showing compact neuroepithelial morphology. Immunostaining reveals PAX6^+^ (green) and Tuj1^+^ (red) neuroepithelial domains. DAPI (blue) marks nuclei. Scale bars: 200 µm. **(b)** H&E and immunofluorescence images of cystic ORG_4 organoids reveal ventricle-like lumens lined with epithelial projections. Immunostaining confirms choroid plexus identity by TTR^+^ (green), ZO-1^+^ (green) epithelial borders and presence of GFAP^+^ (red) astrocytes. Insets highlight apical localization of epithelial markers. DAPI (blue) marks nuclei. Scale bars: 200 µm. **(c)** Quantification of PAX6^+^ and Tuj1^+^ cells in ORG_1 and ORG_2 organoids cultured with or without Matrigel (MG). Matrigel embedding promotes higher Tuj1^+^ cell proportions and lower PAX6^+^ populations. Data are mean ± SEM; ^*^p < 0.05 (unpaired t-test). **(d)** Quantification of TTR^+^ and Tuj1^+^ cells in cystic ORG_4 organoids, showing enrichment for ChP markers. Data are presented as mean ± SEM from n = 6–8 sections per condition.

## 4. Discussion

We present a quick, xeno-free/ serum-free protocol for generating ChP-enriched organoids from human iPSCs using simple combinations of GSK3β inhibition, WNT pathway modulation, and extracellular matrix embedding. This approach reliably produced ventricle-like cystic organoids with hallmark ChP features—including TTR and ZO-1 expression. Notably, we observed the initial appearance of cystic structures as early as day 6, providing an early morphological signature of ChP-like tissue commitment.

Importantly, we demonstrate that early CHIR99021 treatment alone (ORG_3/4) biases toward ChP fate, while the addition of SB505124 (ORG_1/2) steers differentiation toward dorsal forebrain neuroepithelium. These findings highlight the importance of fine-tuned early signalling in directing regional identity in organoid systems. Furthermore, Matrigel embedding not only enhanced organoid size and cyst formation but also promoted neuronal maturation in neuroepithelial organoids, offering a simple route to optimize structural complexity.

Compared to previous ChP organoid protocols requiring long-term culture and multi-step induction, our method is rapid, robust, and accessible, making it suitable for both small-scale disease modelling and high-throughput applications. The early and visible onset of cyst formation adds a valuable morphological readout for real-time monitoring of organoid fate.

While this study focuses on early structural and molecular characterization, future work will explore functional outputs, including CSF-like fluid secretion, barrier integrity assays, and pathogen entry models.

In conclusion, our protocol provides a versatile and efficient platform for ChP development studies, neuroinflammatory disease modeling, and screening of CNS-acting compounds, bridging a gap between neurodevelopmental biology and translational research.

## Acknowledgments

We acknowledge the Ritsumeikan Global Innovation Research Organization (R-GIRO).

## Declarations

### Ethical Approval

Human iPSC line 201B7-Ff employment was approved by the Ritsumeikan University Ethics Committee (No. 2021-004-3)

### Consent to Participate

Not applicable.

### Consent to Publish

Not applicable.

### Authors Contributions

Rodi Kado Abdalkader: Conceptualized and managed the project, designed, and performed experiments, analyzed, and interpreted data, visualized the data, and wrote the manuscript. Takuya Fujita: provided resources and contributed to manuscript refinement. All authors critically reviewed the manuscript and agreed with the publication.

### Availability of data and material

Not applicable.

