## Supplementary Information for "Quick and robust method for the generation of human iPSC-derived choroid plexus organoids"

**Table 1. Chemical reagents list**

| Additives | Working conc. | Target | Maker |
|---|---|---|---|
| IWP-2 | 10 μM | WNT | Santacruz |
| SB-505124 | 10 μM | TGF-β | Santacruz |
| Y-27632 | 10 μM | ROCK | Fujifilm Wako |
| Retinoic acid | 500 nM | RAR | Fujifilm Wako |
| CHIR99021 | 2.5 μM | GSK-3α/β | Santacruz |

**Table 2. Antibodies list**

| Target protein | Host species | Maker | Catalog number | Working dilution |
|---|---|---|---|---|
| PAX6 | Rabbit | Proteintech | 12323-1-AP | 1:500 |
| Beta Tubulin 3/ Tuj1 | Mouse | Gene Tex | GTX27751 | 1:250~ |
| GFAP | Mouse | MBL | D097-3 | 1:100~ |
| TTR | Rabbit | Proteintech | 11891-1-AP | 1:100~ |
| ZO-1 | Rabbit | Proteintech | 21773-1-AP | 1:500~ |
